# TuxNet: A simple interface to process RNA sequencing data and infer gene regulatory networks

**DOI:** 10.1101/337147

**Authors:** Maria Angels de Luis Balaguer, Ryan J. Spurney, Natalie M. Clark, Adam P. Fisher, Rosangela Sozzani

**Affiliations:** Plant and Microbial Biology Department, North Carolina State University, Raleigh, NC 27695; Electrical and Computer Engineering Department, North Carolina State University, Raleigh, NC 27695; Biomathematics Program, North Carolina State University, Raleigh, NC 27695

## Abstract

Predicting gene regulatory networks (GRNs) from gene expression profiles has become a common approach for identifying important biological regulators. Despite the increase in the use of inference methods, existing computational approaches do not integrate RNA-sequencing data analysis, are often not automated, and are restricted to users with bioinformatics and programming backgrounds. To address these limitations, we have developed TuxNet, an integrated user-friendly platform, which, with just a few selections, allows to process raw RNA-sequencing data (using the Tuxedo pipeline) and infer GRNs from these processed data. TuxNet is implemented as a graphical user interface and, using expression data from any organism with an existing reference genome, can mine the regulations among genes either by applying a dynamic Bayesian network inference algorithm, GENIST, or a regression tree-based pipeline that uses spatiotemporal data, RTP-STAR. To illustrate the use of TuxNet while getting insight into the regulatory cascade downstream of the *Arabidopsis* root stem cell regulator *PERIANTHIA (PAN)*, we obtained time course gene expression data of a PAN inducible line and inferred a GRN using GENIST. Using RTP-STAR, we then inferred the network of a PAN secondary downstream gene, *ATHB13*, for which we obtained wildtype and mutant expression profiles. Our case studies feature the versatility of TuxNet to infer networks using different types of gene expression data (i.e time course and steady-state data) as well as how inference networks are used to identify important regulators.

**SUMMARY:** TuxNet offers a simple interface for non-computational biologists to infer GRNs from raw RNA-seq data.

## INTRODUCTION

Knowing whether few or many genes operate together, and whether these genes are involved in unique regulatory circuits, can provide insight into the size and structure of the gene regulatory networks (GRNs) controlling specific biological processes. Currently, gene expression profiling experiments can be used to obtain the compendia of genes expressed either across different cell or tissue types (de Luis Balaguer et al. 2017; S. Li et al. 2016), developmental time (Brady et al. 2007), or in response to distinct stimuli (Seki et al. 2002). Accordingly, RNA-sequencing (RNA-seq) methods are used in combination with differential gene expression analyses to investigate gene expression changes in different cell- and tissue-types or conditions, such as a wild type versus mutants or control versus treatments. Different analytical pipelines are used to process RNA-seq reads, obtain quantified gene expression (normalized gene counts), and perform statistical tests to identify differentially expressed genes (DEGs). Bioinformatics tools performing these analyses include Bowtie2 (Ben Langmead and Salzberg 2012), BWA (H. Li and Durbin 2009), HTSeq-count (Anders, Pyl, and Huber 2015), baySeq (Hardcastle and Kelly 2010), DESeq (Hardcastle and Kelly 2010), DESeq2 (Love, Huber, and Anders 2014), edgeR (Robinson, McCarthy, and Smyth 2010), and Tuxedo (TopHat + Cufflinks + Cuffdiff) (Trapnell et al. 2012). The resulting list of DEGs are often used by downstream analytical tools, such as GENIST (de Luis Balaguer et al. 2017), GENIE3 (Huynh-Thu et al. 2010), and ARACNE (Margolin et al. 2006), which infer causal relationships among DEGs to, for example, identify key regulators. Despite the importance of these analyses, current tools for both processing RNA-seq data and inferring GRNs are not integrated and are often not automated, requiring users to have bioinformatics and/or programming acquaintance (Trapnell et al. 2012; Robinson, McCarthy, and Smyth 2010).

To provide an integrated tool for processing sequencing data as well as inferring GRNs, we have developed TuxNet, a Matlab graphical user interface (GUI) that can be used with minimal programming and bioinformatics knowledge. We show that TuxNet, sequentially: 1) identifies DEGs from next generation sequencing data using fastq-mcf (Aronesty 2011; Aronesty 2013), Tuxedo (Trapnell et al. 2012), and TuxOP (a custom-built Matlab pipeline that processes the Tuxedo pipeline outputs); and 2) infers, from spatiotemporal data, GRNs using either GENIST (de Luis Balaguer et al. 2017) or a regression tree-based pipeline (RTP-STAR) (Huynh-Thu et al. 2010; Shibata et al. 2018). Through a case study aimed to obtain insight into the regulatory cascade downstream of a key *Arabidopsis* root stem cell regulator, *PERIANTHIA (PAN)* (de Luis Balaguer et al. 2017), we show how TuxNet takes raw sequencing data as input files and returns different output files, including tables with normalized gene expression values, lists of DEGs, and, using different types of gene expression data, inferred causal relationships among genes. Specifically, time course gene expression data of a PAN inducible line were used to infer a network using GENIST, and wildtype and mutant expression profiles of a PAN downstream gene, *ATHB13*, were used to infer a GRN using RTP-STAR. Importantly, through our examples, we guide the user on how to select the few parameters needed to run the GUI, which we provide as an open source package (https://github.com/madeluis/TuxNet).

## RESULTS

### Overview of the three tabs of TuxNet

TuxNet provides a user-friendly environment where users can automatically process gene expression data (RNA-seq data), identify differentially expressed genes (DEGs), and infer gene regulatory networks (GRNs) from the processed data. TuxNet is developed as a graphical user interface (GUI) divided in three tabs, TUX, GENIST, and RTP-STAR (Fig. 1, Fig. 2, and Fig. 3). The three TuxNet tabs have consistent file formatting to ensure that analyses, when expanding more than one tab, do not require data manipulation or coding by the user.

**Fig. 1.**
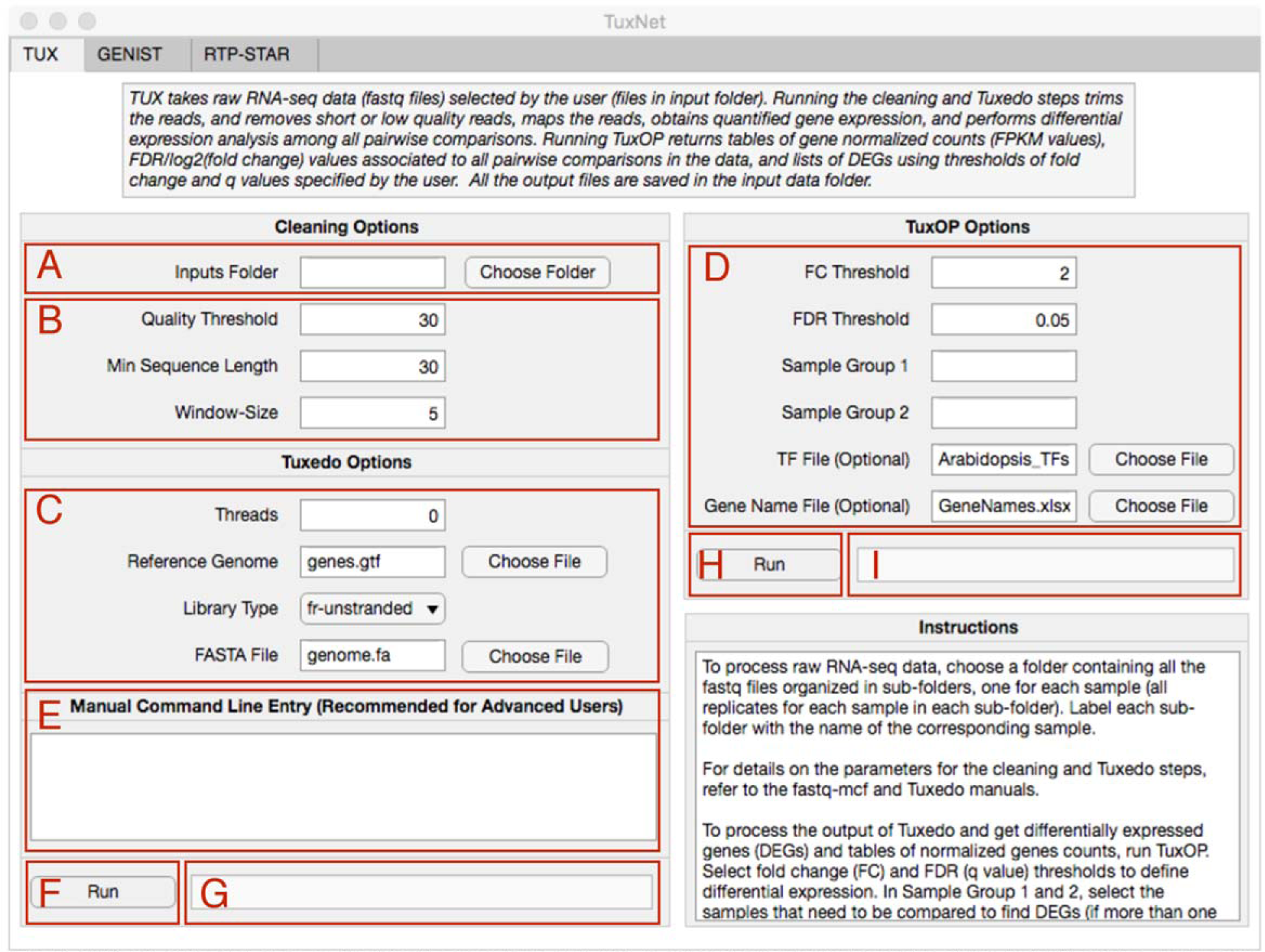
TUX tab of TuxNet shown with default parameters. **A** Button to select the folder that contains all the fastq files to be analyzed in the TUX tab. **B** Parameters to run the cleaning step through fastq-mcf. Values shown are default parameters. **C** Parameters to run the Tuxedo pipeline. Values shown are default parameters. **D** Parameters to run TuxOP (processing of the Tuxedo pipeline output). Values shown are default parameters to find differentially expressed genes (thresholds for differential expression –fold change or FC, and FDR or q values–and samples to compare during the differential expression test). **E** Text box to run fastq-mcf or the Tuxedo pipeline using command line syntax with custom parameters. **F** Button to run the fastq files processing steps (fastq-mcf and the Tuxedo pipeline). The button will run either the data and parameters inserted through the text boxes and drop down menus, or the manual command line entry. **G** Error message output. **H** Button to run TuxOp. **I** Error message output.

**Fig. 2.**
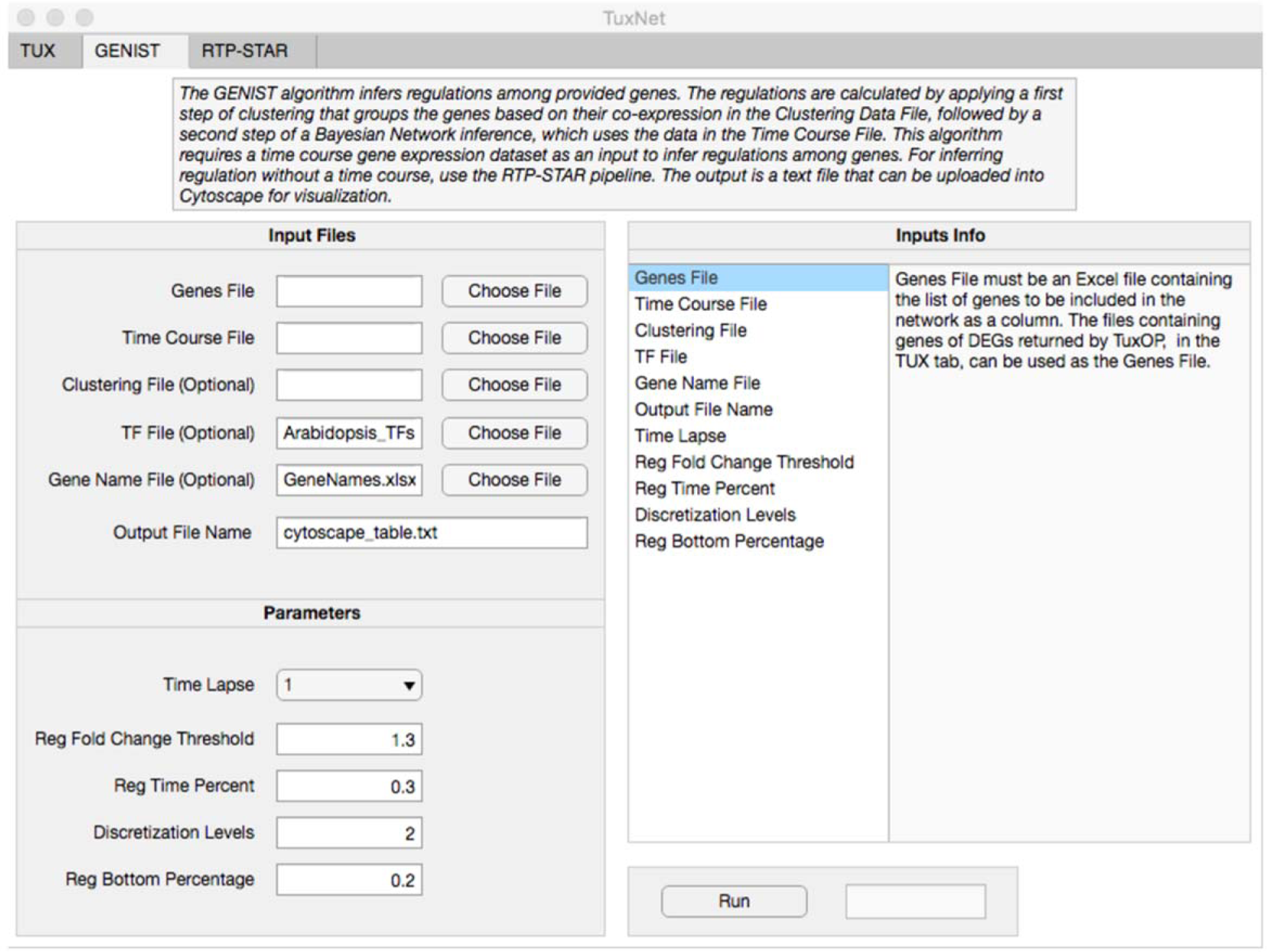
GENIST tab of TuxNet shown with default parameters. **A** Buttons to select the required input files. **B** Buttons to select the optional input files. **C** Text file to type in a name for the output table that will contain all the inferred network regulations generated by GENIST. Name shown is the default. **D** Parameters to run GENIST. Values shown are default parameters. **E** Information textbox to provide explanations on each of the input files and parameters. **F** Button to run GENIST after all files and parameters have been selected. **G** Error message output.

**Fig. 3.**
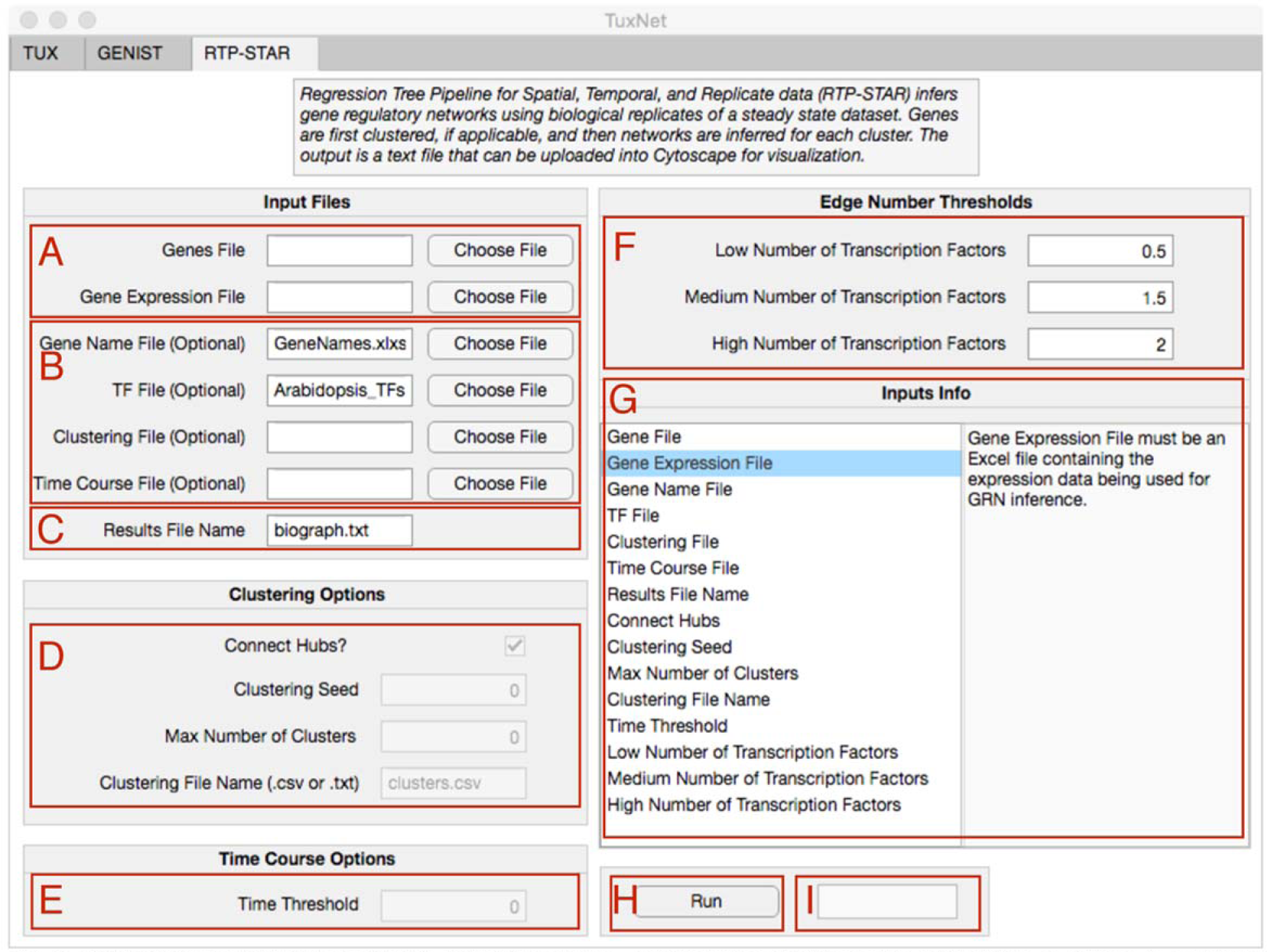
RTP-STAR tab of TuxNet shown with default parameters. **A** Buttons to select the required input files. **B** Buttons to select the optional input files. **C** Text file to type in a name for the output table that will contain all the inferred network regulations generated by RTP-STAR. Name shown is the default. **D** Parameters to run RTP-STAR associated to the clustering step. They get active if a clustering file is chosen as an input in B. **E** Parameter to run RTP-STAR associated to the time course input file (step for inferring sign of regulations). It becomes active if a time course file is chosen as an input in B. **F** Default parameters to set the number of edges to be kept for a low (first value), medium (second value), or high (third value) number of transcription factors. **G** Information textbox to provide explanations on each of the input files and parameters. **H** Button to run RTP-STAR after all files and parameters have been selected. **I** Error message output.

The TUX tab allows users to process RNA-seq data in the form of fastq files, as usually returned by an Illumina sequencer. Accordingly, the TUX tab, which implements fastq-mcf (Aronesty 2011; Aronesty 2013) and the Tuxedo pipeline (TopHat + Cufflinks + Cuffdiff) (Trapnell et al. 2012), performs cleaning, mapping, count, and normalization of the RNA-seq raw reads, as well as differential expression analysis. The RNA-seq processing workflow (fastq-mcf + Tuxedo) runs after uploading the raw RNA-seq data and selecting only a few parameters. To simplify the user interface, we offer the possibility of specifying, through individual text boxes, only a few parameters for fastq-mcf and Tuxedo. However, any additional parameters available to run these packages can be specified by typing a command line to call the executable files (fastq-mcf, TopHat, Cufflinks, Cuffmerge and Cuffdiff) in the text box as shown in Supplemental Fig. 1.

To start the analysis of the RNA-seq reads, the user imports a folder containing the fastq files (raw RNA-seq data) obtained as a result of an experimental design (e.g., a treatment and a control or a time-course experiment) (Fig. 1A). Note that this input data folder is where all the files resulting from the analysis, including the intermediate steps, will be saved. If the RNA-seq experiment is designed to encompass multiple samples (biological conditions), such as wild-type versus mutant or control versus treatment, then the folder must be organized in sub-folders, each containing all the fastq files (all biological replicates) corresponding to each sample. TUX processes these fastq files using fastq-mcf (ea-utils) to remove the Illuminan adapters (barcodes) and eliminate the short and low quality reads (the TuxNet package incorporates a file, IlluminaAdaptorSeq.fasta, which contains the adapters that will be trimmed) (Fig. 1B). The resulting clean fastq files are saved in the input data folder as clean_*.fastq, where * refers to each of the input fastq file names. TUX uses TopHat (Trapnell, Pachter, and Salzberg 2009) to take the clean fastq files, map the reads to the reference genome, and return a *_thout folder associated to each input fastq file, that includes .bam files of mapped reads. By implementing Cufflinks (Roberts et al. 2011), TUX uses the .bam files to assemble the transcripts and estimate their abundances and, associated to each input fastq file, returns a *_clout folder with the assemblies. TUX uses Cuffmerge to merge the Cufflinks assemblies together and returns its results in the merge_out folder. By using Cuffdiff, TUX uses the mapped reads and the assemblies to find significant changes in transcripts and gene expression, and returns an output folder, cuffdiff_out, containing multiple files with transcript and gene abundance as well as splicing information. Importantly, contained in cuffdiff_out, the gene_exp.diff file provides gene-level pairwise comparison information between all the samples processed, such as normalized read counts in Fragments Per Kilobase of transcript per Million mapped reads (FPKM) units, and false discovery rate (FDR) values (Fig. 1C). Although to run the described Tuxedo steps the user must select the reference genome and indexed FASTA files associated to each organism, we include an Arabidopsis reference genome and indexed FASTA file in the TuxNet package as the default option of the GUI. If the user needs to use a different organism, or a different reference genome, the reference genome and indexed FASTA files can be selected in the “Reference genome” and “FASTA file” select menus (Fig. 1C). Moreover, although the GUI does not offer the option of selecting a reference transcriptome, the user can run the Tuxedo pipeline with a reference transcriptome by typing the command lines to call the executable files (Supplemental Fig. 1., Fig. 1E) following directions as in (Trapnell et al. 2012). Similarly, analyzing paired-end reads is not supported through the TUX automated options, thus fastq files containing paired-end reads should be analyzed through the command line entry box (Supplemental Fig. 1., Fig. 1E). In our examples throughout the next section, we provide parameter values that can be chosen by the user.

In addition to fastq-mcf and Tuxedo, the TUX tab implements TuxOP, which we designed to run after the fastq-mcf and Tuxedo pipelines and allow users to find DEGs in either pairwise or combinatorial sample comparisons. Specifically, TuxOP processes gene_exp.diff, returns a table containing mean gene expression values as well as gene expression values for each biological replicate (Fig. 1D). In addition, TuxOP provides a table containing a list of all the DEGs, where differential expression is defined by user-specified FDR and fold change thresholds. The list of DEGs obtained after comparing gene and transcript expression across the different samples are saved as DEG_**vs***.xlsx, where ** and *** indicate the names of the samples that are compared. Optionally, the user can import a transcription factor (TF) list (see Materials and Methods, Formats of Input Files) to allow TuxOP to return, in addition to the DEGs, a list of the TFs contained among the DEGs. TuxNet includes a complete TF list for Arabidopsis that is pre-selected by default, but users can select a different file if a different organism is used, or, alternatively, select no file. TuxOP also allows the user to select a gene name file (see Materials and Methods, Formats of Input Files) to name the DEGs. As before, TuxNet includes an Arabidopsis gene name file that is pre-selected by default, but users can select a different file (to work with a different organism or with customized names) or, again, select no file (to set all gene names as the gene identifier).

GENIST and RTP-STAR, the second and third TuxNet tabs, are used to infer GRNs from the list of DEGs and the expression tables returned by TUX. However, the GENIST and RTP-STAR tabs are not dependent on the TUX tab and users can infer GRNs with expression data that has been processed with other bioinformatics pipelines, such as edgeR, or DESeq, and formatted as specified in Formats of Input Files. To run GENIST, the user imports an .xlsx file with a list of genes to be included in the network (gene file, Fig. 2A). Since GENIST is a Dynamic Bayesian Network (DBN)-based inference algorithm that infers gene-gene regulations when time-course gene expression data are available (de Luis Balaguer et al. 2017), the user also selects a .xlsx file, such as the table gene_expression.xlsx returned by TuxOP, with the time-course experiment that will be used to infer the network (see Materials and Methods, Formats of Input Files) (Fig. 2A). The DBN will provide information about the probability of gene-gene relationships with any time-course data, such as time series taken after, for example, a treatment or an induction, or a developmental progression of a cell line and/or tissue. The time course can contain evenly or unevenly spaced time points to accommodate the requirements of the biological experiment. However, to permit the calculation of probabilities, the data set should have a minimum of 3 time points. If only the gene and the time course expression files are provided, GENIST infers a GRN based on the expression of the user-specified genes throughout the time-course experiment.

Clustering based on gene expression can lead to groups of genes that are spatially co-expressed or functionally related. Clustering genes before the DBN inference step has been shown to improve GENIST performance (de Luis Balaguer et al. 2017). This step reduces the complexity of the inference step and becomes critical for inferring large networks. Therefore, TuxNet allows the addition of a clustering step prior the network inference in GENIST, which we recommend applying whenever clustering data are available. The clustering data could be either a cell- or tissue-type, a mutant or overexpressor, or another time course expression dataset. To run the clustering step before the inference then, in addition to the gene file and the expression data, the user imports an .xlsx file with the expression data that will be used for clustering (see Materials and Methods, Formats of Input Files) (Fig. 2B). Note that the datasets used for clustering should not be used for the inference.

GENIST offers the option of allowing only TFs to be regulators in the inferred network. To choose this option, the user imports a TF list (see Materials and Methods, Formats of Input Files) (Fig. 2B). Again, TuxNet includes a complete TF list for Arabidopsis that is pre-selected by default, but users can select a different file. In addition, GENIST allows the user to select a gene name file (see Materials and Methods, Formats of Input Files) (Fig. 2B) that will be used to name the genes in the final network. Also in this case, TuxNet includes an Arabidopsis gene name file that is pre-selected by default that users can change by selecting a different file. With the Genes File, the Time Course File, and any of the described optional files, GENIST infers a DBN and returns a graphical representation of the network. In addition, GENIST returns a .txt file containing all the edges of the network, i.e., source gene, target gene, type of regulation (+/−1, for activation/repression, or 0 for undetermined), and weight (associated to the inferred probability of each regulation) of each edge. If a clustering file is provided, in addition to the final network, GENIST returns a graphical representation of the sub-networks inferred for each cluster and a .txt file indicating which genes are members of each cluster number. The table containing the final network can then be imported into a network visualization software, such as Cytoscape (Shannon et al. 2003), to obtain representations of the network with higher quality graphics and customizable options.

RTP-STAR implements a regression tree algorithm, GENIE3 (Huynh-Thu et al. 2010), and uses the biological replicates of steady-state (i.e., one time point) gene expression experiments to infer GRNs. To run RTP-STAR, the user imports an .xlsx file with a list of genes that will be included in the network (gene file, Fig. 3A) (see Materials and Methods, Formats of Input Files). The user also imports an .xlsx file, such as the table gene_FPKM_replicates.xlsx returned by TuxOP, with the expression data of all the biological replicates of all samples that will be used to infer the network (see Input Files)(Fig. 3A). With these input files, RTP-STAR infers a GRN by running a regression tree pipeline on the biological replicate data for the genes of interest (Huynh-Thu et al. 2010; Shibata et al. 2018). After the inference step, RTP-STAR trims the number of edges in the network to keep only the edges with the highest confidence (the edges that are most likely to be true positives). The edges are trimmed according to the ratio of transcription factors to genes, as datasets with a higher number of TFs should have more potential for genetic regulation than datasets with a low number of TFs. As detailed for GENIST, RTP-STAR allows for the addition of a clustering step prior to the network inference to reduce the complexity of the inference step (Fig. 3B). If clustering data are included, RTP-STAR clusters the genes before performing inference. RTP-STAR also allows the user to import a TF list, to permit only TFs to be regulators, as well as a gene name file that will be used to name the genes in the network (Fig. 3B). Moreover, RTP-STAR offers the possibility of including an additional temporal dataset to infer the type of the predicted regulations (activation, repression, or undetermined) (Materials and methods) (Fig. 3B). If a time-course dataset is not included, RTP-STAR assumes all the inferred regulations in the network have an undetermined sign. With these inputs, RTP-STAR returns a graphical representation of the network (one graph for each cluster, if clustering is applied), as well as a .txt file containing all the regulations of the inferred network, i.e., source gene, target gene, and type of regulation (activation, repression, or undetermined) of each edge. As before, this table can then be imported into network visualization software tools, such as Cytoscape (Shannon et al. 2003), to obtain customizable representations of the network.

Overall, TuxNet can be run by importing the data and selecting a few options and parameters. By allowing the users to select their processed gene expression files to run GENIST and RTP-STAR, we offer versatility for inferring GRNs with data analyzed with any bioinformatics pipeline in addition to Tuxedo. Similarly, we offer pre-selected Arabidopsis files to automatically run all the TuxNet tabs, but offer flexibility to choose other files, consequently permitting the use of TuxNet with any organism.

### Network inferences from time course transcriptomic data using GENIST

To guide the user through the execution of the TUX and GENIST tabs of TuxNet, here we provide a case study that shows how to infer a network with GENIST using an RNA-seq time-course gene expression dataset processed with TUX. For this, we obtained a time course of the Arabidopsis root at times 0, 6, and 24 hours after *PERIANTHIA (PAN)* induction using the XVE:PAN TRANSPLANTA line (Materials and Methods, de Luis Balaguer et al. 2017), which can be used to identify genes downstream of *PAN* and infer gene regulations. Moreover, since we are inferring a network downstream of PAN and *PAN* is specifically expressed in the quiescent center (QC) (de Luis Balaguer et al. 2017), we used a QC specific gene expression dataset obtained from sorting and profiling pWOX5::erGFP in wildtype and *pan* mutant backgrounds (de Luis Balaguer et al. 2017) for the clustering step. We then combined the QC-specific and the *PAN* time course datasets to find DEGs downstream of *PAN* specifically in the QC. We used these DEGs to infer our network using the Dynamic Bayesian Network algorithm used by the GENIST tab. Below we provide details to run this analysis with TuxNet.

To process the time-course data and find DEGs using TUX, we selected the folder containing the input files, which, in our example, consists of nine fastq files (three replicates of each of the three time points, T0i.fastq.gz, T1i.fastq.gz T2i.fastq.gz, for i=1,2,3) (Fig. 4A). Importantly, before running TuxNet, we subdivided the folder in three subfolders, each containing the three biological replicates of each time point. After importing the fastq files, we selected the parameters for running the cleaning step (to remove adapter sequences, low quality reads, and short reads) as well as the parameters for running the Tuxedo pipeline as shown in Box 1 and Fig. 4B. Running the TUX tab generates output files from each of the intermediate steps, as shown in Supplemental Fig. 2, which are saved in the input data folder. In the last step of the Tuxedo pipeline implemented in the TUX tab, a test of differential expression between each pairwise comparison within all the processed samples is performed. This test returns, among others, gene-level log2 fold changes and FDR values (q values) for all the pairwise comparisons. In our example, the pairwise comparisons are T0-T1, T0-T2, and T1-T2 (./cuffdiff_out/gene_exp.diff).

**Fig. 4.**
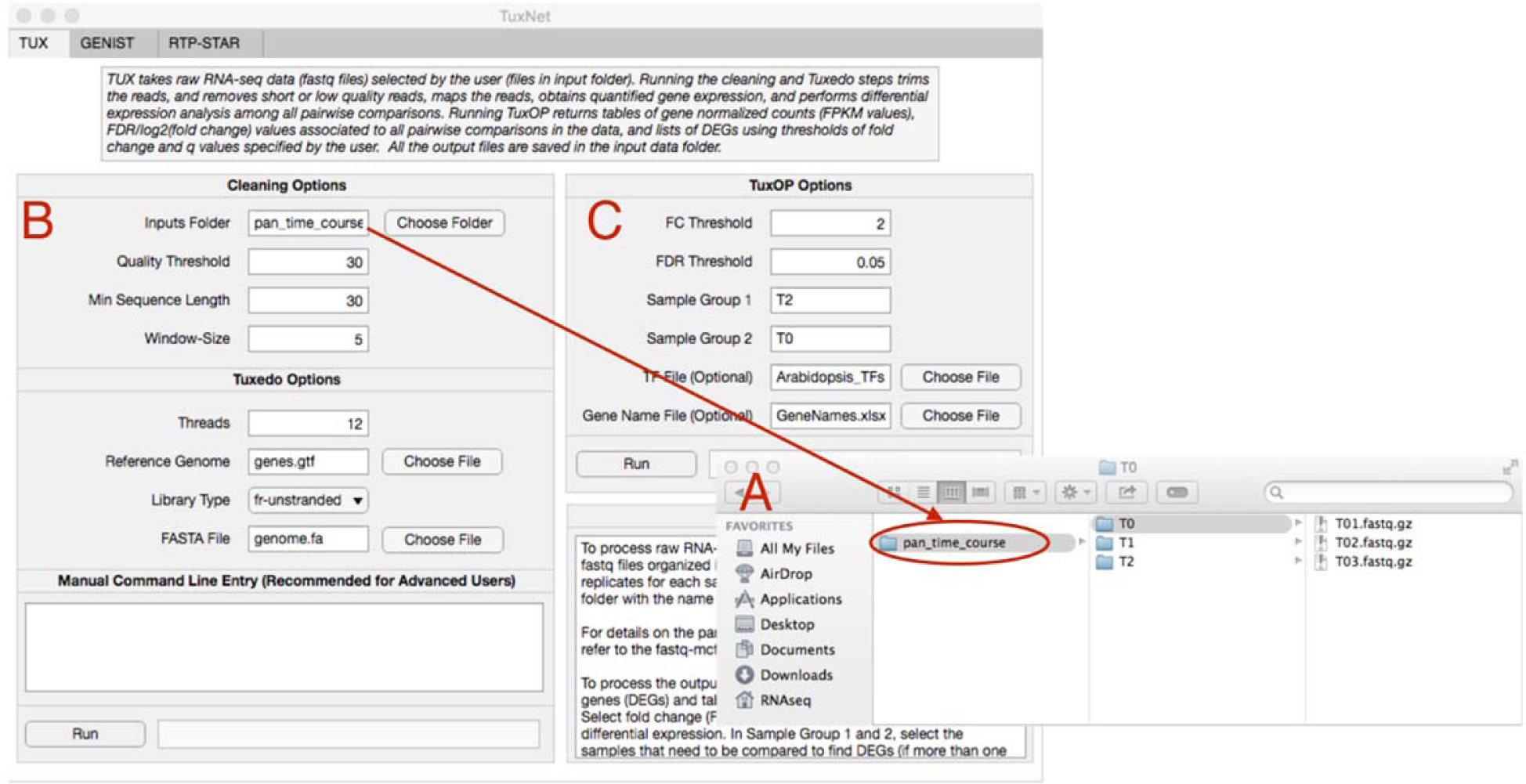
TUX tab of TuxNet shown with input files and parameters used to run the PAN network illustrated in Fig. 6. **A** Window showing the folder that is selected, containing the input files (fastq files), to run the pipeline. The folder is organized in sub-folders, each one containing all the biological replicates for each sample. **B** Parameters selected to run fastq-mcf and the Tuxedo pipeline on the input files. **C** Parameters selected to run TuxOP on the output of the Tuxedo pipeline to find DEGs. The names provided in Sample Group 1 and Sample Group 2 match the names of the sub-folders that are imported as an input (T0,T1,T2).

To generate user-customized tables of DEGs and tables of gene expression values that can be used as input files by GENIST and RTP-STAR, the user can then run TuxOP, which we designed to specifically provide flexibility in the identification of DEGs. Accordingly, TuxOP processes the output returned by Tuxedo (gene_exp.diff), performs an automatic search of DEGs (not only in pairwise comparisons, but also in group comparisons, such as ((T0 & T1) vs T2), and returns the tables of DEGs and expression values. TuxOP allows the users to choose which combination of samples they want to compare to obtain lists of DEGs, and which thresholds of fold change and q values they prefer for identifying differential expressed genes. To obtain different lists of DEGs, TuxOP can be run multiple times using different sample combinations. TuxOP becomes particularly useful when searching for DEGs in group comparisons from large datasets. In our example, with only three samples (T0, T1, and T2), there are 9 different pairwise and group comparisons (T1 vs T0, T2 vs T0, T2 vs T1, (T1 & T2) vs T0, etc). If, in addition, the user is interested in identifying only genes up- or down-regulated in each comparison (up-regulated in (T1 & T2) vs T0, down-regulated in (T1 & T2) vs T0, etc.), the number of possible comparisons becomes 18. Since this number increases exponentially with the number of samples contained in the dataset, the automation to identify DEGs from combinatorial comparisons is therefore very advantageous.

In our example, we show this specific feature by identifying two individual lists of genes that are either activated or repressed by PAN after 24 hrs. For this, we focused on genes that were differentially expressed at T2 vs T0 after PAN induction. To find these genes, we ran TuxOP with the parameters we have listed in Box 2, where we set the thresholds for differential expression as q<0.05 and fold change (FC) > 2. We ran TuxOP with the value T2 in Sample group 1 and with the value T0 in Sample group 2, to obtain a list of DEGs in T2 vs T0 (Box 2, Fig. 4C). Note that we can run TuxOP with the value T0 in Sample group 1 and with the value T2 in Sample group 2, which leads to the same results. If the users are interested in identifying genes that are differentially expressed in T1 vs T0, as well as in T2 vs T0, they can specify T1, T2 in Sample group 1 and T0 in Sample group 2. Similarly, users can specify T0 in Sample group 1 and T1, T2 in Sample group 2. Moreover, users interested in obtaining the expression tables, but not in finding lists of DEGs, can run TuxOP by selecting any value in Sample group 1 and Sample group 2. Running TuxOP with the set of parameters specified in Box2 generates two files, DEG_T0_vs_T2.xlsx (Dataset 1), containing the list of DEGs up-regulated in T0 vs T2, and DEG_T2_vs_T0.xlsx (Dataset 2), containing the list of DEGs up-regulated in T2 vs T0. The first sheet of each excel file lists all the DEGs, and the second sheet of the excel file lists only the TFs that were differentially expressed. In addition, TuxOP generates a table that contains a summary of the ./diff_out/gene_exp.diff file, complete_table.xlsx (expression values of all genes in all samples, q values in all comparisons, and fold changes in all comparisons). TuxOP also returns a table of the mean expression values for each gene in all the samples processed as gene_expression.xlsx (Dataset 3), as well as a table of the gene expression values for each of the replicates processed as FPKM_replicates.xlsx. All tables generated by TuxOP are saved in the folder of the input files and can be directly used to run the GRN inference pipelines within TuxNet. We next analyzed the RNA-seq profiles obtained from QC cells in WT and *pan* mutant to select DEGs, as shown in Box 1. As the analysis of this dataset had been previously published (de Luis Balaguer et al. 2017), we selected previously used parameters to find DEGs, i.e., q < 0.05, FC > 1. We chose Sample group 1 = pan, Sample group 2 = WT to obtain a list of 3397 DEGs (and 274 TFs) (upregulated in the *pan* mutant, DEG_pan_vs_WT.xlsx, Dataset 4, and downregulated in the *pan* mutant, DEG_WT_vs_pan.xlsx, Dataset 5) (note that the same results are obtained by opting for the selection: Sample group 1 = WT, Sample group 2 = pan). In addition to the DEGs, TuxOP returned a table of the mean expression values for each gene from all the processed samples (gene_expression.xlsx, Dataset 6). To find the set of genes downstream of *PAN* to include in the network, we then manually intersected the lists of genes found to be downstream of *PAN* in the mutant and in the inducible line (803 genes, Dataset 7).

Given that the precision of the inferred edges decreases as 1) the number of genes included in the network increases, and 2) the number of time points used to infer the edges decreases, users can choose to infer a network with TFs only to reduce the complexity of the GRN. Since we obtained a large number of DEGs from the intersection (803 genes) and our time-course dataset contained only three time points, we chose to infer a TF regulatory network (TFRN). Therefore, we specifically intersected the TFs upregulated in the mutant and downregulated in the inducible line at T2, and the TFs downregulated in the mutant and upregulated in the inducible line at T2 ((DEG_pan_vs_WT.xlsx ∩ DEG_T0_vs_T2.xlsx) ∪ (DEG_WT_vs_pan.xlsx ∩ DEG_T2_vs_T0.xlsx)). The final list of TFs used to infer the network is listed as DETF_pan.xlsx (Dataset 8). After generating all the necessary gene files, using the parameters as listed in Box 3, we ran GENIST with the *pan* mutant data for clustering and the *PAN* inducible time course data for inferring regulations (edges) (Box 3, Fig. 5, Supplemental Table 1). This resulted in a table containing all the inferred regulations, saved in the folder of input files as cytoscape_table_pan.txt (Dataset 9), and a graphical representation of the network. Note that the genes included in the final table are only those that have, at least, one connecting edge.

**Fig. 5.**
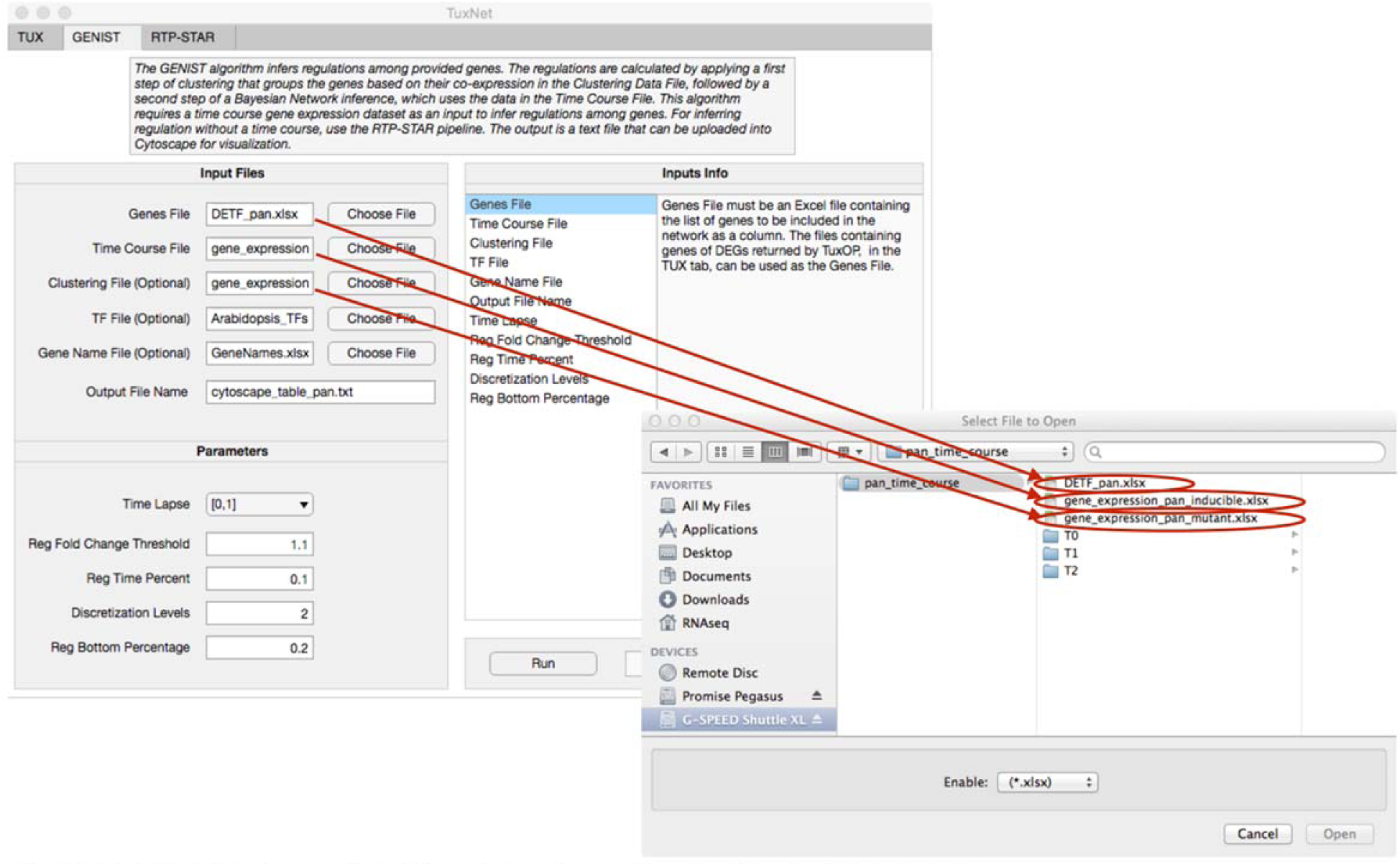
GENIST tab of TuxNet shown with input files and parameters used to run the PAN network illustrated in Fig. 6.

The file returned by GENIST can be imported into Cytoscape to obtain customized graphical representations of the network, as shown in Fig. 6. In our example, we chose to plot the size of the nodes as a function of the number of genes that they directly regulate. These customized plotting features allow users to visually identify network characteristics. Specifically, by varying node sizes, we aimed to highlight the network’s main regulators. In the resulting network (Fig. 6), as expected, *PAN* is shown as the gene with the largest number of regulations (outgoing edges). In agreement with previous studies (de Luis Balaguer et al. 2017), the network shows *PAN* regulating key genes involved in various root stem cell functions. Accordingly, out of the 64 TFs in the *PAN* network, 19 TFs, including (*PLETHORA1*) *PLT1*, (*AINTEGUMENTA-LIKE6*) *AIL6*, and (*BABY BOOM*) *BBM*, were all previously shown to be downstream of *PAN* and have a role in stem cell specification (de Luis Balaguer et al. 2017). Additionally, the network representation suggests that 29 of the remaining factors, which include genes such as *IAA16, JAZ4*, and *ATHB13*, with no known previous role in stem cell function, are part of a secondary regulatory cascade. Thus, to test whether these genes may have a role in stem cell regulation as well as to test the functionality of TuxNet with steady-state data, we examined the network of a *PAN* secondary downstream gene.

**Fig. 6.**
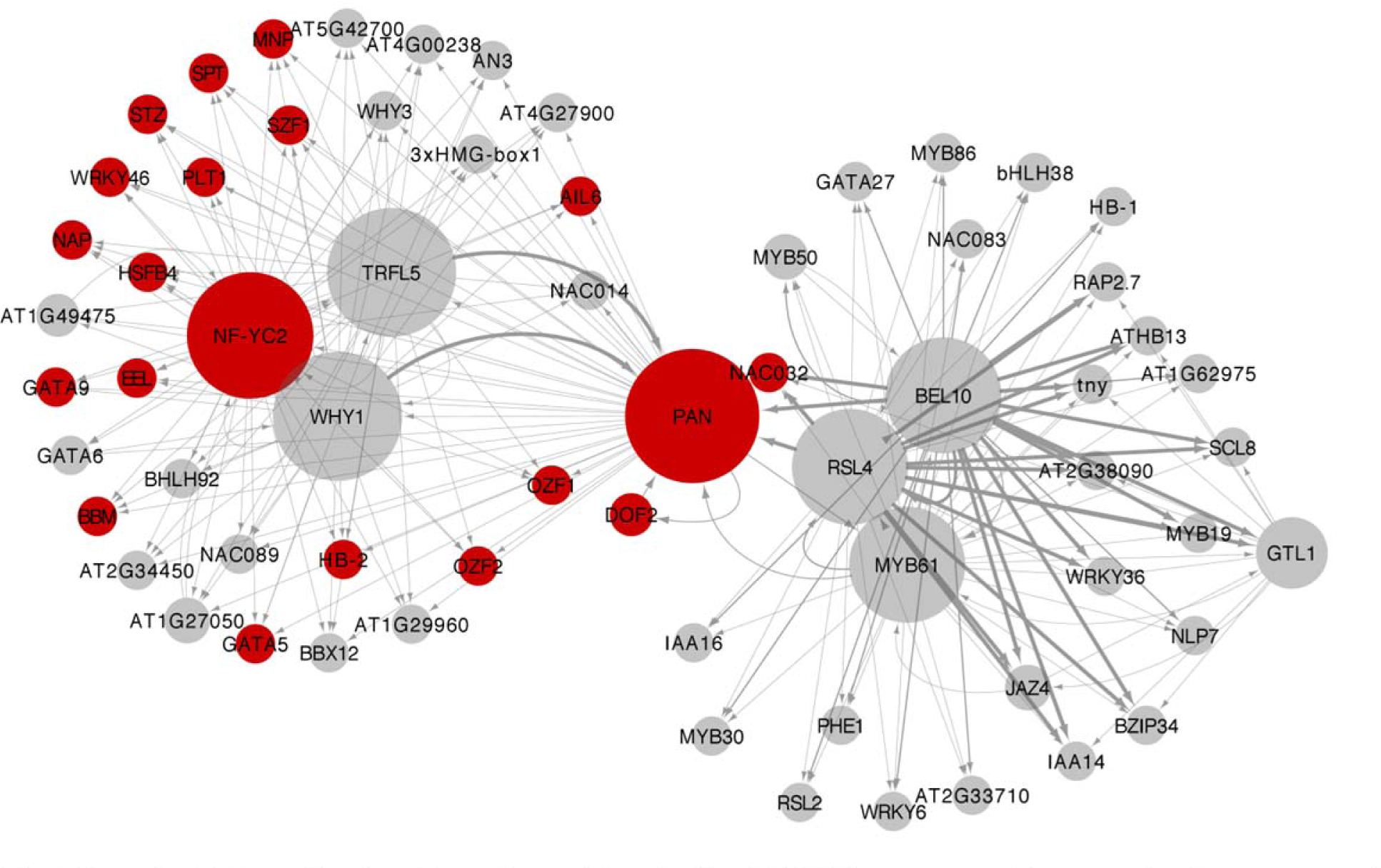
Network of *PAN* and its downstream factors inferred with GENIST. The structure of the network reflects a temporal aspect of the *PAN* regulatory cascade: a first regulatory cascade of *PAN* (left), and a secondary cascade, through the *MYB61* node (right). The size of the nodes correlates with the number of genes that they regulate. Nodes in red indicate genes previously shown to have a role in stem cell regulation. The shape of the arrow on each edge indicates whether the predicted regulation is an activation 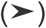, a repression (**|**), or it has an undetermined sign (•).

### Network inferences from biological replicates of transcriptomic data using RTP-STAR

Here we provide a case study that shows how to infer a network with RTP-STAR using an RNA-seq steady-state gene expression dataset processed with TUX. For this, we obtained three biological replicates of RNA-seq from *athb13* mutant and wild type (WT) control roots (Materials and Methods). Accordingly, *ATHB13* was among the genes in the secondary network of *PAN*, and *athb13* mutant alleles showed a disorganized stem cell niche, suggesting a role of *ATHB13* in stem cell regulation (Supplemental Fig. 7). To infer a network that reflected the regulatory cascade from *PAN* to *ATHB13* and their downstream genes, we identified DEGs downstream of *ATHB13*, and next, we manually combined the DEGs downstream of *PAN* and *ATHB13*. Additionally, since RTP-STAR allows the user to combine biological replicates from different experiments, we combined the gene expression values from each biological replicate from the *PAN* time-course dataset analysis and the *ATHB13* dataset analysis to build a dataset for inferring regulations among the DEGs. As for GENIST, we provide the example and rationale for using a clustering dataset prior to the inference step. Specifically, for clustering, we used the transcriptional profile obtained from root tissues as in (S. Li et al. 2016), since the expression data used for inferring the network comprise whole-root expression profiles. Below we provide details to run this analysis with TuxNet.

**Fig. 7.**
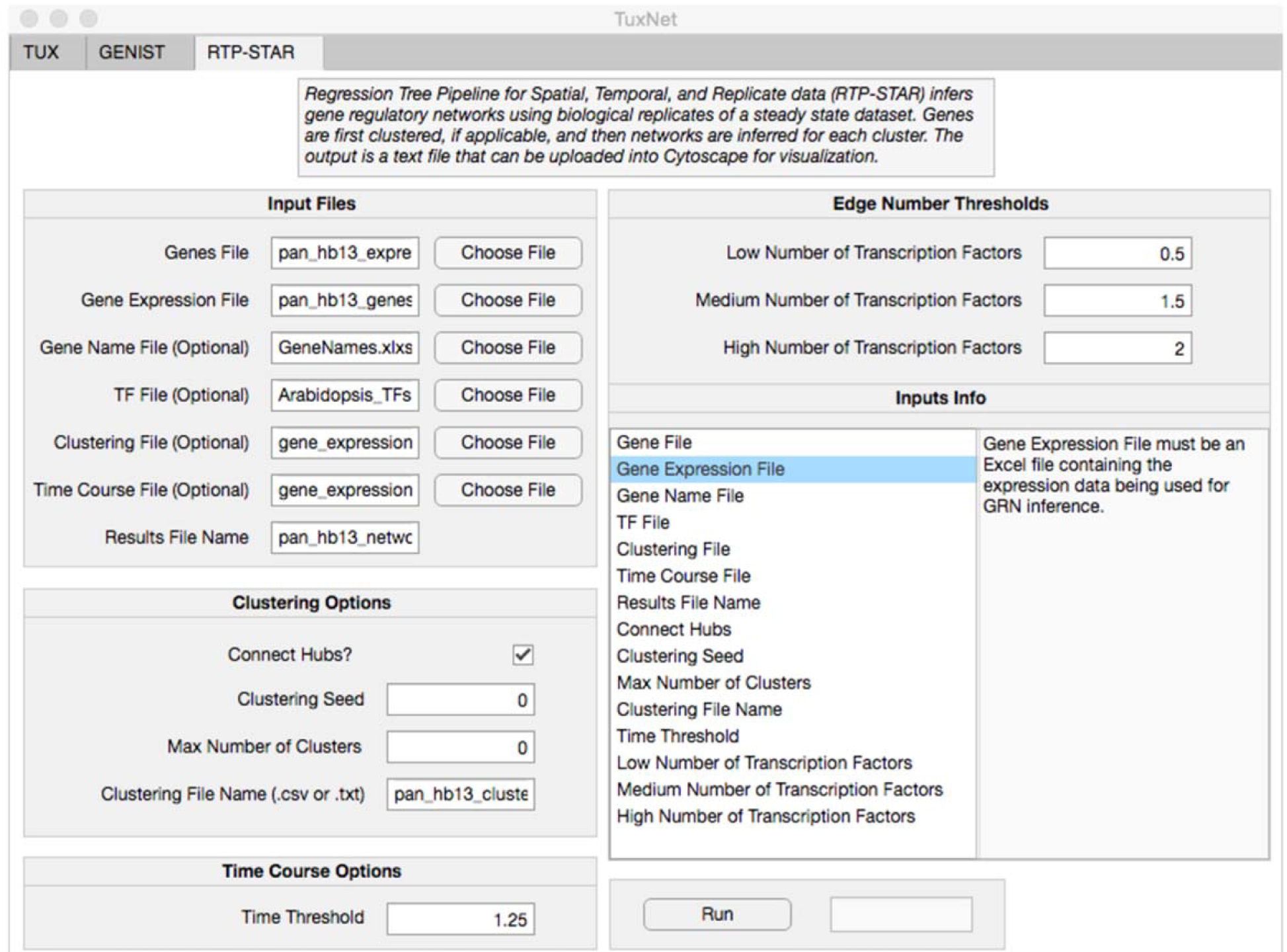
RTP-STAR tab of TuxNet shown with input files and parameters used to run the *PAN-ATHB13* network shown in Fig. 8. 0 values indicate default values.

To process the WT and *athb13* mutant gene expression profiles, we analyzed all the sequencing files (three replicates of *athb13* mutant roots and WT roots), using TUX, following the steps as listed in Box 1. The file FPKM_replicates.xlsx (Dataset 10), returned by the TUX analysis, contains the FPKM values of all the biological replicates processed in the dataset. Importantly, this is the file containing the gene expression values used by RTP-STAR to infer the networks. In our case, FPKM_replicates.xlsx contains the normalized expression values for the three *athb13* replicates and the three WT replicates. In addition, the TUX analysis (with q<0.05 and FC > 2, as shown in Box 2), returns two files of DEGs (DEG_ATHB13_vs_WT.xlsx and DEG_WT_vs_ATHB13.xlsx). The genes in these two files together resulted in 481 DEGs, up- or down-regulated in the mutant, between *athb13* and WT (Dataset 11). To obtain the final list of genes to be included in the network, we manually combined the list of these 481 DEGs with the 65 TFs from the *PAN* analysis (DEG_ATHB13_vs_WT.xlsx U DEG_WT_vs_ATHB13.xlsx U DETF_pan.xlsx) (Dataset 8) resulting in a total of 544 genes (pan_hb13_genes.xlsx) (Dataset 12). To obtain the combined *athb13* mutant*-PAN* time course expression dataset, we merged the biological replicates of *athb13* mutant and WT roots (FPKM_replicates.xlsx resulting from the analysis of the *athb13* and WT dataset), with the biological replicates of the *PAN* inducible line at T0 and T2 (FPKM_replicates.xlsx resulting from the analysis of the PAN inducible line at T0 and T2). We used the resulting merged table as the expression data (Dataset 13). Given the large number of genes, we clustered the genes before the inference step. As before, we analyzed the clustering dataset (NCBI SRA database, BioProject PRJNA323955) (S. Li et al. 2016) using TuxNet (Box 1), and selected the gene_expression.xlsx returned by TuxOP as the Clustering File (Dataset 14). Finally, to predict which regulations are activations or repressions, we used a time-course dataset to infer the sign of the regulations in the network. Since *PAN* is expressed in the root stem cells (de Luis Balaguer et al. 2017) and *athb13* mutant showed a disorganized stem cell niche, the time-course dataset we used consisted of five time points from the Arabidopsis root stem cells, every 24 hours, from days 3 to 7 (de Luis Balaguer et al. 2017). We analyzed these data as in Box 1 and Box 2 using TuxNet, and used the resulting gene_expression.xlsx as the time-course data to infer the sign of the regulations (Dataset 15). We then inferred the *ATHB13* GRN using RTP-STAR using the input files and parameters specified in Box 4 (Fig. 7). Running RTP-STAR with these parameters results in a graphical representation of the network (divided in one graph per cluster) as well as the pan_hb13_network.txt (Dataset 16) file, which contains a table with all the regulations inferred in the final network. Note that the genes included in the final table are only those that have at least one connecting edge. We then imported the file returned by RTP-STAR into Cytoscape to obtain a customized graphical representation of the network, as shown by the Cytoscape plot of pan_hb13_network.txt (Fig. 8). As described for GENIST, we chose to vary the size of the nodes as a function of the number of genes that they directly regulate to highlight the network’s main regulators. Accordingly, *PAN* is found at the top of the network regulating 151 genes through direct and indirect edges, while *ATHB13* regulates a subset of 116 genes.

**Fig. 8.**
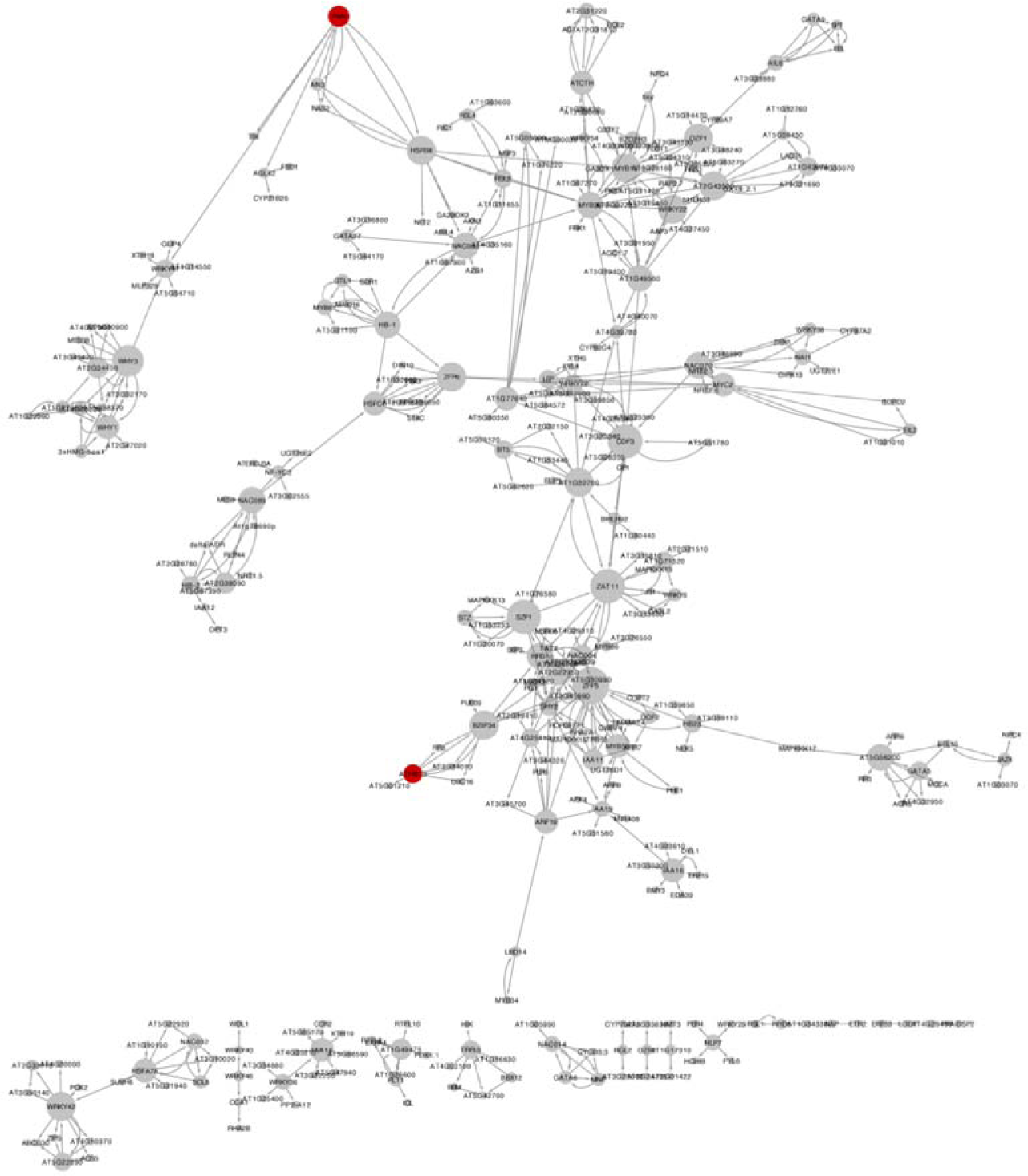
Network of *PAN, ATHB13*, and their downstream factors, inferred with RTP-STAR. The size of the nodes correlates with the number of genes that they regulate. The shape of the arrow on each edge indicates whether the predicted regulation is an activation 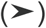, a repression (**|**), or it has an undetermined sign (•).

Here, while getting insight into the regulatory cascade downstream of the *Arabidopsis* root stem cell regulator *PERIANTHIA (PAN)*, we show how TuxNet can help users process RNA-seq data using a GUI instead of requiring users to write and run code using a programming language. Accordingly, we show that while all the tabs of TuxNet require users to set a few (<6) parameters, we provide default values. Importantly, we provide parameter information and range values (Supplemental Table 1, 2 and Supplemental Fig. 3-6) to guide users in the selection of alternative parameter values. We show how users can infer a GRN using the GENIST tab when experimentally time course gene expression data are acquired or RTP-STAR with steady-state data. Overall, we illustrate the versatility of TuxNet to infer networks with many different types of gene expression data and how the networks can be used to identify important regulators in a regulatory cascade.

## DISCUSSION

In this manuscript, we present TuxNet, a graphical user interface (GUI) specifically designed for non-computational biologists with minimal programming and bioinformatics skills. TuxNet allows, by selecting just a few parameters from text boxes and dropdown menus, to process user-specified gene expression data and, importantly, infer GRNs from the processed data. Through our examples, we provide critical information on how to select each of the parameters needed to run the three tabs of the GUI, namely TUX, GENIST, and RTP-STAR. We offer flexibility to process gene expression data from any organism with TuxNet by incorporating the option to select a reference genome (genes.gtf) and an indexed FASTA file (genome.fa) in the TUX tab. As a default configuration, we provide these files for Arabidopsis, as large amounts of gene expression data are obtained from this model organism. However, if data from other organisms need to be processed, we allow users to simply import these organism-specific files into, for example, the TuxNet directory, and select them from the TUX tab. Moreover, as part of the TUX tab, we have designed TuxOP, which allows users to specify differential expression criteria and obtain “on-demand” lists of DEGs with their corresponding false discovery rate (FDR) and fold-change thresholds from either pairwise or combinatorial comparisons. Importantly, all the tables and intermediate data resulting from each step are used as inputs for the next step. Thus, as opposed to current network inference tools, TuxNet integrates all the steps of the analysis from the data processing to the network inference. Among all bioinformatics pipelines that can be used to perform this RNA-seq data analysis, we had opted for fastq-mcf + Tuxedo because these can be fully run from the computer terminal, and therefore, can be run from the Matlab terminal and packaged into our Matlab-based GUI (Materials and Methods). Since the GRN inference tabs, GENIST and RTP-STAR, have no dependences on the data processing step, users can process their RNA-seq data using any bioinformatics pipelines in addition to Tuxedo. However, to infer causal relationships among DEGs, users should select GENIST when a time-course gene expression dataset is available, or RTP-STAR, when biological replicates of a steady state dataset is available.

Although our aim is to provide case studies to show the utility of TuxNet, by generating gene expression data, we also gained insight into the regulatory cascade downstream of a key *Arabidopsis* root stem cell regulator, *PAN.* Specifically, to infer a network with GENIST, we obtained an RNA-seq time-course gene expression dataset of the Arabidopsis root after *PAN* induction at times 0, 6, and 24 hours. Since the time-points were unevenly spaced out, we chose a value of 0 and 1 for the GENIST parameter “time-lapse “, to explore the possibility that factors regulate genes within the same time point (which could more likely occur when the experimental design has longer time steps, i.e 12 hrs), or with a time lag of 1 time step (which could occur when time steps are short, i.e 30min). The resulting inferred network suggests that *PAN* regulates genes in two regulatory cascades. First, PAN regulates a group of 35 TFs that contain 18 factors that had been previously shown to be stem cell specific. Among these are important stem cell regulators, such as (*PLETHORA1*) *PLT1*, (*AINTEGUMENTA-LIKE6*) *AIL6*, and (*BABY BOOM*) *BBM*. In a secondary cascade (“secondary network “), PAN is predicted to regulate 29 factors that, with the exception of *NAC032* (de Luis Balaguer et al. 2017), had not been shown to be specifically expressed in the stem cells. To gain more insight into these genes, we examined the “secondary network” of *PAN*. We specifically inspected *ATHB13*, for which we observed a disorganized stem cell niche in its mutant alleles. In the network incorporating all the DEGs downstream of *PAN* and *ATHB13, PAN* is still at the top of the inferred regulatory cascade, while *ATHB13* regulates a smaller sub-network in a feedback loop with *BZIP34,* suggesting a less important role of ATHB13 in regulating stem cell genes.

We believe that our presented tool, which incorporates previously validated algorithms for network inference, can be of great use to scientists interested in not only processing their RNA-seq data but also performing downstream analyses to further mine their data. We foresee that the balance between automation and flexibility in our tool will allow it to be applied to multiple organisms and be used by scientists with a variety of computational backgrounds.

## ACKNOWLEDGEMENT

We thank Jose Alonso, Anna Stepanova, Bob Franks, and Terri Long, of North Carolina State University, for critically reading the manuscript. Support for this work was provided by the National Science Foundation (NSF) (CAREER MCB 1453130) and bilaterally by the NSF and the Biotechnology and Biological Sciences Research Council (BBSRC) (NSF MCB-1517058) (to R.S.); Funding to N.M.C. is provided by NSF (DGE-1252376); Funding to A.P.F. is provided by NSF (DGE-1252376).

## METHODS

### TuxNet architecture

The GUI is implemented as a Matlab app that calls each of the scripts, or executables, running each task of the workflow. Specifcally, Fastq-mcf and Tuxedo run from the executables provided in (Aronesty 2011; Aronesty 2013; Trapnell et al. 2012). TuxOP runs from Matlab scripts specifically developed for this GUI: find_DEG.m, create_Exp_Table.m, create_DE_Table.m, fpkm_replicates_init.m, and fpkm_replicates.m. GENIST fully runs from Matlab scripts we previously published in (de Luis Balaguer et al. 2017). RTP-STAR also fully runs in Matlab and implements GENIE3 as explained in (Huynh-Thu et al. 2010; Shibata et al. 2018). Specifically developed for this GUI, RTP-STAR provides the possibility of clustering before inference and/or using time course data to predict the type of regulation (activation or repression). Additionally, RTP-STAR trims the number of edges returned by GENIE3 based on the ratio of transcription factors to genes and infers the type of regulation (activation, repression, or undetermined) using the same first-order Markov assumption as GENIST (de Luis Balaguer et al. 2017). The TuxNet app offers additional tools, which include Sam tools (H. Li et al. 2009) and Bowtie software (B Langmead et al. 2009).

### Running requirements

The use of TuxNet requires a 64-bit computer running Mac OS X (10.4 Tiger or later), with Matlab installed (R2017b Matlab version or later). GENIE3, available from http://www.montefiore.ulg.ac.be/∼huynh-thu/GENIE3.html, must be also installed before RTP-STAR can be run using TuxNet. To install GENIE3, download the MATLAB .zip folder, unzip the files, and follow the installation instructions included in the documentation. TuxNet can be downloaded from https://github.com/madeluis/TuxNet. Download all the files in the TuxNet project as a .zip folder. To open and run TuxNet, open the TuxNet.mlapp file, included in the folder. To run TuxNet for the first time using Arabidopsis data, unzip genome.fa.zip (included in the folder).

### Formats of input files

#### gene file

.xlsx file with all the genes provided in the first column of the first worksheet. The first row of the sheet is considered a header and will not be processed. The sheet can contain additional columns but only the first column is read by TuxNet.

#### *time course expression file* (GENIST only)

.xlsx file, such as the table gene_expression.xlsx returned by TuxOP, containing the time-course expression data that will be used to infer the network. This file contains the gene names in the first column of the first sheet, and can contain all genes in the genome (the genes to be included in the networks do not need to be pre-sorted).

#### *time course expression file* (RTP-STAR only)

.xlsx file, such as the table gene_expression.xlsx returned by TuxOP, containing the time-course expression data that will be used to infer the sign of the regulations. This file contains the gene names in the first column of the first sheet, and can contain all genes in the genome (TuxNet searches for the genes from “gene file” in this file).

#### *expression file* (RTP-STAR only)

.xlsx file, such as the table FKPM_replicates.xlsx returned by TuxOP, containing the biological replicate expression data that will be used to infer the network. This file contains the gene names in the first column of the first sheet, and can contain all genes in the genome (TuxNet searches for the genes from “gene file” in this file.)

#### clustering file

.xlsx file with all the genes provided in the first column of the first worksheet. The first row of the sheet is considered a header and will not be processed. The sheet can contain additional columns but only the first column will be imported. The file can contain all genes in the genome (TuxNet searches for the genes from “gene file” in this file).

#### TF file

.xlsx file containing the gene identifiers of the known TFs in the first column of the first sheet.

#### gene name file

.xlsx file containing a list of characterized genes with their names. The first and second column of the first sheet should contain the locos ID of these genes and the gene names, respectively.

### Plant Material and Growth Conditions

*Arabidopsis thaliana* Columbia (Col-0) seeds were used. The XVE:PAN transgenic line under the control of the β-estradiol-inducible promoter was previously described in (Coego et al. 2014; de Luis Balaguer et al. 2017). Induction of *PAN* expression was completed as described in (Coego et al. 2014). The *athb13-1* and *athb13-2* mutant seeds were previously described in (Ribone, Capella, and Chan 2015).

XVE:PAN and *athb13-*1 seeds were wet sterilized using 50% bleach, 100% ethanol, and rinsed 6 times with water. After sterilization, the seeds were stratified and imbibed at 4 degrees Celcius for 2 days. The seeds were plated on 1X Murashige and Skoog (MS) medium supplemented with 1% sucrose on top of Nitex mesh and grown vertically at 22 degrees Celsius in long-day conditions (16-hours light/ 8-hours dark). *athb13* seeds were grown for 5 days on 1X MS supplemented with 1% sucrose. XVE:PAN seeds were first grown on 1X MS supplemented with 1% sucrose plates, and then were transferred to 1X MS supplemented with 1% sucrose and 10 uM β-estradiol for 0, 6, or 24 hour *PAN* induction. All XVE:PAN seeds were grown for a total of 5 days.

### Microscopy and Phenotypic Analysis

Phenotypic analysis of WT and *athb13* mutants was completed using the ZEISS LSM 710 confocal microscope. The 488nm laser was used for red channel acquisition. A 10 uM propidium iodide solution was used to stain the cell walls for visualization. Images were taken with ZEN software (Zeiss).

### *athb13* Transcriptional Profile and *PAN* Inducible Line Time Course

For the *PAN* inducible line time-course experiments at 0, 6, and 24 hours after induction as well as WT and *athb13* transcriptional profiles, roots were dissected at approximately 0.5 cm from the root tip. RNA was extracted using the RNeasy Micro Kit (Qiagen). cDNA synthesis, amplification, and library preparation were performed using the NEBNext Poly(A) mRNA Magnetic Isolation Module and the NEBNext Ultra II Directional RNA Library Prep Kit for Illumina (NEB). Libraries were sequenced on an Illumina HiSeq 2500 with 100 bp single-end reads.

### Accession numbers

The data reported in this paper have been deposited in the Gene Ex-pression Omnibus (GEO) database, https://www.ncbi.nlm.nih.gov/geo (accession numbers GSE112563, and GSE112564

##### BOX1

Selection of the TUX tab parameters used to analyze the *PAN* inducible line time course (see user manual of fastq-mcf (Aronesty 2011; Aronesty 2013) and the Tuxedo pipeline (Trapnell et al. 2012) for parameter information):

1. Set the quality threshold. All bases below the threshold will be removed. We set this parameter to 30, which indicates that the bases kept have 99.9% accuracy.
2. Set a window size for quality trimming. If no window size is selected (equivalent of window size = 1), quality trimming is performed by scanning the read, one base pair at a time, and clipping the read if a base pair that does not meet the quality threshold is encountered. If a window size w > 1 (for example, w = 3pb to 6bp) is selected, clipping will occur if the average quality score of w consecutive base pairs falls below the quality threshold. Here, we set this parameter to 5bp. If on average, 5 base pairs have a quality score lower than the quality threshold, the read is clipped from that position. In addition to performing quality trimming, trimming is applied to remove the Illumina adapters (as specified in IlluminaAdaptorSeq.fasta) without the need to specifying additional parameters.
3. Set a minimum read length. All reads below this length will be removed. This parameter can be set close to the read length of the RNA-seq data (e.g 70bp-80bp for raw RNA-seq reads of 100bp). To allow shorter clipped reads to be kept, the minimum read length can be set lower. We set this parameter to 30bp.
4. Select the reference genome. We used the pre-selected option, which is the Arabidopsis gtf file (genes.gtf).
5. Select the Arabidopsis indexed FASTA file. We used the pre-selected option, which is the Arabidopsis indexed FASTA file (genome.fa).
6. Set the number of threads for running the pipeline. This parameter should be less or equal to the number of cores of the computer used for the analysis. We set this parameter to 12 (12-core computer used).
7. Specify the type of library used to generate the samples. The options are fr-unstranded, fr-firststrand, and fr-secondstrand. We set this parameter as fr-unstranded, according to our library preparation.

##### BOX2

To obtain the list of DEGs from the *PAN* induction time course shown in Datasets 1 and 2, we run TuxOP with the following parameters:

1. FC threshold = 2. This sets the log2 fold change of expression cutoff for differential gene expression.
2. FDR threshold = 0.05. This sets the FDR (q value) cutoff for differential gene expression.
3. Gene name File: Locus_Primary_Gene_Symbol_2013.xlsx (default, pre-selected file).
4. TF list file: Arabidopsis_TF file.xlsx (default, pre-selected file).
5. Sample group 1 = T2, Sample group 2 = T0, to obtain DEGs at time T2 vs T0.

##### BOX3

To infer the *PAN* network, the GENIST tab requires the selection of the following parameters (see Input File Formats for information on the formatting of the input tables, and Supplemental Table 1 and Supplemental Fig. 3-6 for parameter definitions):

1. Genes File: DETF_pan.xlsx, generated after intersecting the DETFs in the *pan* mutant and inducible line (Dataset 8).
2. Time Course File: gene_expression.xlsx returned by TuxOP from the time-course data analysis (Dataset 3).
3. Clustering File: gene_expression.xlsx returned by TuxOP from the *pan* mutant data analysis (Dataset 6).
4. Gene name File: Locus_Primary_Gene_Symbol_2013.xlsx (default, pre-selected file).
5. TF list file: Arabidopsis_TF file.xlsx (default, pre-selected file).
6. Time Lapse: [0 and 1]. We offer the possibility to select either no time lapse [0, which assumes A regulates B within the same time point] or a time lapse [1, which assumes A regulates B at the next time point] (see Supplemental Fig. 3 for parameter explanations). Given our time course (0, 6, and 24hrs) we selected 0 and 1 as we assume that the regulation may happen within and between time points.
7. Number of discretization levels of the expression values: 2 (default). To calculate the Bayesian probabilities we selected two discretization levels, each gene is considered to have either a high or a low expression at each time point (Supplemental Fig. 4).
8. Regulators Fold Change Threshold: 1.1. By setting this value, only TFs whose expression changes at least 10% before a change of expression of their target are considered as regulators (Supplemental Fig. 5).
9. Regulators Time Percent: 0.1. By selecting this value, only TFs showing a change of expression during, at least 10% of the time points of the time course, are considered as regulators (Supplemental Fig. 6).
10. Regulations Bottom Percentage: 0.2. This value filters out the bottom 20% of the regulations.
11. Output file: cytoscape_table_pan.txt

##### BOX4

To infer the *PAN* and *ATHB13* network, the RTP-STAR Tab requires the selection of the following parameters (see Input File Formats for formatting of the input tables, and Supplemental table 2 and Supplemental Fig. 5 for parameter definitions):

1. Genes File: pan_hb13_genes.xlsx (Dataset 12).
2. Gene Expression File: pan_hb13_exprssion.xlsx. This file combines the FPKM_replicates.xlsx returned by TuxOP from the *ATHB13* data analysis with the replicates of T0 and T2 from the FPKM_replicates.xlsx returned by TuxOP from the *PAN* time course data analysis (Dataset 13).
3. Clustering File: gene_expression.xlsx returned by TuxOP from the analysis of the BioProject PRJNA323955 dataset (Dataset 14).
4. Gene name File: Locus_Primary_Gene_Symbol_2013.xlsx (default, pre-selected file).
5. Time course File: gene_expression.xlsx returned by TuxOP from the time time-course dataset (5 time points) from the Arabidopsis root stem cells (Dataset 15).
6. TF file: Arabidopsis_TF file.xlsx (default, pre-selected file).
7. Results file name: pan_hb13_network.txt (default name is biograph.txt), name of the file to save the final table containing all the network regulations to.
8. Connect hubs: true (default value), performs an inference step to connect the hubs of each cluster
9. Clustering seed: N/A (default value), allows the clustering seed to be set so that the clusters are the same every time. The seed is a number that initializes the random number generation. Therefore, changing the seed will change how the genes are clustered in the initial step. If the clustering seed is N/A, the seed will change every time, and could result in different clusters each time the pipeline is run.
10. Max number of clusters: N/A (default value is floor(number of genes/10)+5), where floor() is the closest integer value (for example, floor(11.2) = 11, and floor (13.7) = 13)), allows the user to set the maximum number of clusters. Note that the minimum number of clusters is determined based on the maximum number.
11. Clustering file name: pan_hb13_clusters.txt (default name is clusters.csv), name of the file to save the clusters to.
12. Time threshold: 1.25 (default value), determines genes that change significantly overtime. A gene should change 25% between time points, otherwise those time points are not used to calculate the sign of the regulation (see Supplemental Fig. 5 for parameter explanations).
13. Edge number: N/A (default value is 0.5 for low, 1.5 for medium, and 2 for high number of TFs), sets the number of edges to keep for a low (<1/4 genes are TFs), medium (1/4<number of TFs<1/2), and high (>1/2 genes are TFs) number of TFs. The number of edges kept is then calculated as edgenumber*number of genes

**Author contributions**
M.A.L.B. designed the experiments, developed scripts integrated in the GUI, analyzed the data, and wrote the paper; R.J.S. developed the interface; N.M.C. developed scripts integrated in the GUI and complemented the writing; A.P.F. performed the experimental work; R.S. designed and supervised the experiments, and complemented the writing.

